# Fibronectin-mediated inflammatory signaling through integrin α5 in vascular remodeling

**DOI:** 10.1101/2021.01.06.425577

**Authors:** Madhusudhan Budatha, Jiasheng Zhang, Martin A. Schwartz

## Abstract

Adhesion of vascular endothelial cells (ECs) to the underlying basement membrane potently modulates EC inflammatory activation. The normal basement membrane proteins laminin and collagen IV attenuate inflammatory signaling in part through integrin α2β1. In contrast, fibronectin, the provisional matrix protein found in injured, remodeling or inflamed vessels, sensitizes ECs to inflammatory stimuli through integrins α5β1and and αvβ3. A chimeric integrin in which the cytoplasmic domain of α5 is replaced with that of α2 pairs with β1 and binds fibronectin but signals like α2β1. Here, we examined mice in which integrin α5 is replaced with the α5/2 chimera, using the transverse aortic constriction (TAC) and partial carotid ligation (PCL) models of vessel remodeling. Following TAC and PCL surgery, WT mice showed increased fibronectin deposition and expression of inflammatory markers, which were strongly attenuated in a5/2 mice. α5/2 mice also showed reduced artery wall hypertrophy in the TAC model and diminished inward remodeling in the PCL model. Acute atherosclerosis after PCL in hyperlipidemic ApoE−/− mice on a high fat diet was dramatically decreased in α5/2 mice. These results underscore the key role for integrin α5 signaling in pathological vascular remodeling and support its potential as a therapeutic target.

## Introduction

Arteries remodel in response to physiological stresses such as changes in blood pressure or fluid shear stress, and to pathological stresses such as low and disturbed shear stress, hyperlipidemia, diabetes or oxidative stress among others ^1, 2^. Physiological remodeling is characterized by homeostasis, in which key variables return to their original levels or set point after a perturbation ^1, 3, 4^. For example, increases in blood pressure induce hypertrophic thickening of the vessel wall to restore tensile wall stress to close to original values. Increased or decreased blood flow through a vessel with concomitant changes in fluid shear stress results in outward or inward remodeling of vessel diameters to restore shear stress to close to original values. Inflammation is an essential component of all remodeling ^5^. Cells of the vascular wall that register changes in homeostatic variables (wall or fluid shear stress) activate inflammatory pathways, resulting in recruitment of leukocytes, most prominently monocytes. These cells contribute to these processes by extracellular matrix (ECM) production and degradation, and cytokine/chemokine secretion. Restoration of wall or fluid shear stress to homeostatic levels leads to suppression of inflammatory signaling and restoration of the normal state.

Pathological remodeling, exemplified by atherosclerotic lesions, resembles physiological remodeling in many respects but differs in that the initiating stimuli are never terminated and the inflamed state never resolved ^4, 6^. But ECM remodeling is a central element of both physiological and pathological processes ^6, 7^. Vascular remodeling is universally accompanied by expression and assembly of provisional ECM proteins such as fibronectin (FN), thrombospondin or fibrin, whereas vessel stabilization and quiescence are associated with loss of these components and assembly of basement membranes where collagen IV (coll IV) and laminin (lam) are the main protein components ^8, 9^. Cells interact with provisional ECM proteins through RGD-binding integrins, of which integrins α5β1 and αvβ3 are the most prominent, whereas RGD-independent integrins α2β1, α6β1 and α6β4 are the main receptors for collagens and laminins ^10^.

Importantly, ECM remodeling also modulates inflammatory activation. In endothelial cells, lam/coll IV basement membranes limit inflammatory responses through signaling by integrin α2β1, while FN enhances responses to inflammatory stimuli, including disturbed fluid shear stress, IL1-b and oxidized LDL through integrin α5β1 and αvβ3 ^9^. Though less studied, coll/lam basement membranes also promote smooth muscle cell (SMC) quiescence and differentiation ^11, 12^. The effects of these different ECM proteins were traced to the integrin α subunit cytoplasmic domains: a chimeric integrin in which the α5 cytoplasmic domain was replaced with that of α2 binds fibronectin and supports cell adhesion and cytoskeletal organization normally but signals as if the cells were on lam or coll ^12^. A mouse in which this mutation was inserted into the α5 locus developed normally and is healthy and fertile but showed reduced endothelial inflammatory marker expression in regions of disturbed flow, developed smaller and less inflamed atherosclerotic lesions in hyperlipidemia, and showed improved recovery from hindlimb ischemia ^12, 13^.

In the current study, we analyzed integrin α5/2 mice in three new models of vascular remodeling. We report that these mice show dramatic decreases in inflammation and remodeling, both physiological and pathological.

## Results

### Transverse aortic constriction (TAC)

To assess the role of integrin α5 signaling in pressure-induced artery remodeling, we subjected mice to TAC or to sham surgery as a control. This was accomplished by placing a clamp between the innominate artery and the left carotid, a well-established experimental method to increase pressure in the right carotid artery ^14–16^ (Figure 1A). Left and right carotid arteries were collected one week post-surgery. We observed significant increases in vessel area and lumen diameter of the right carotid artery compared with sham, or with the contralateral left carotid (Figure 1B,C). These changes were accompanied by strongly elevated fibronectin deposition in the endothelial layers of right carotids but not in contralateral left carotids and sham carotids (Figure 1D).

**Figure 1:**
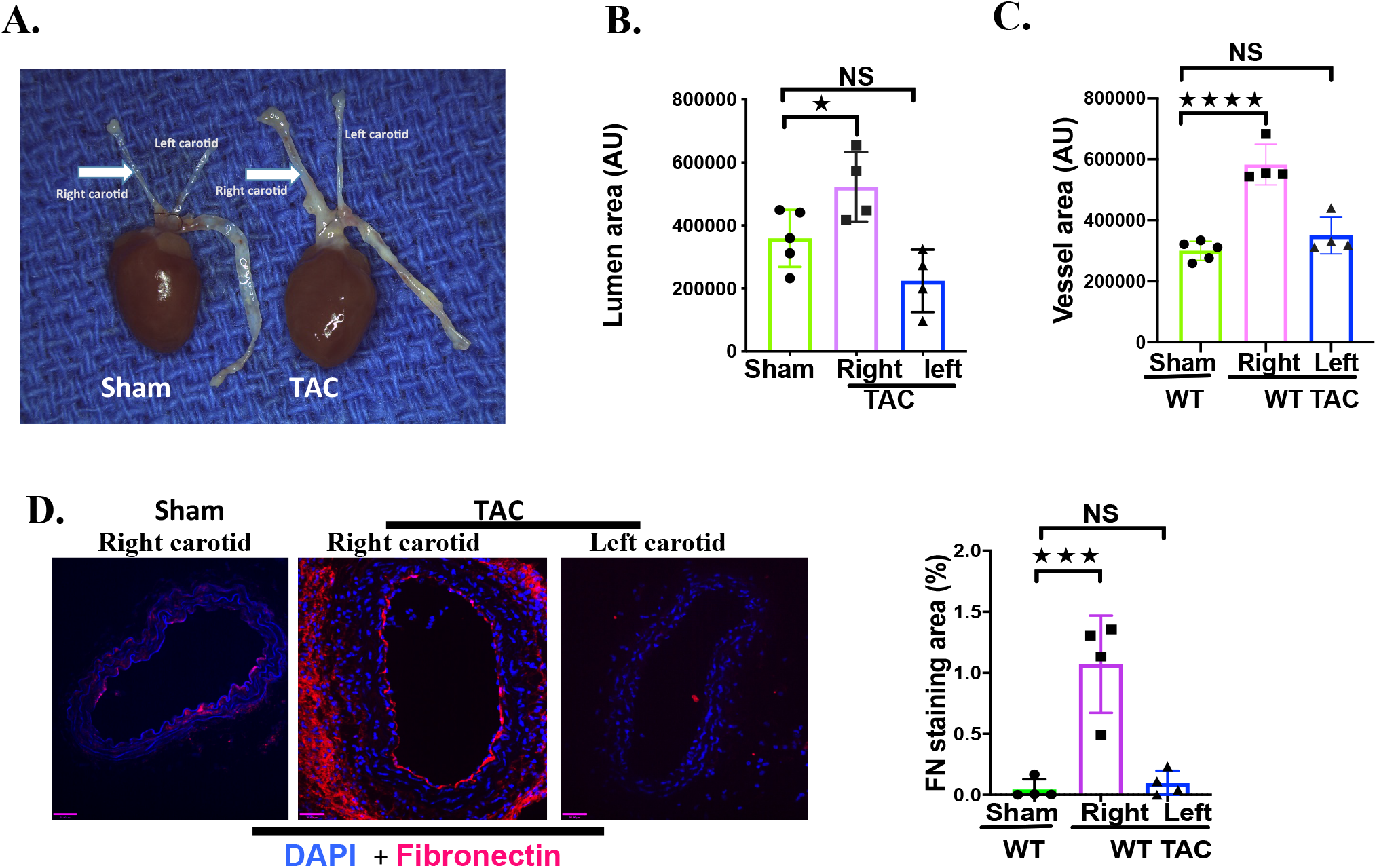
Transverse aortic constriction-induced artery remodeling and fibronectin deposition: **A.** WT mice subjected to sham and TAC surgery. Macroscopic observation revealed thickening of the right carotid artery compared to left and sham carotid arteries**. B & C.** Morphometric analysis revealed a significant increase in the vessel area and lumen diameter of the right carotid artery compared with sham or with contralateral left carotids. **D.** TAC induced deposition of the fibronectin in right carotid artery compared to sham and left carotid. Nuclei were counter stained with DAPI (blue). WT sham mice, n=5; WT TAC surgery mice, n=4; Statistical analysis: one-way ANOVA with Tukey’s post hoc analysis; values are means ± SEM (*p<0.05, ***p<0.001, ****p<0.0001, NS, not significant, compared with WT sham.

### Transverse aortic constriction in α5/2 mice

To investigate the role of fibronectin-integrin α5 signaling in TAC-induced structural remodeling of the carotid arteries, we compared these results to integrin α5/2 knock-in mice. Right carotids from α5/2 knock◻in mice enlarged less than WT mice (Figure 2A,B). The accumulation of FN seen in WT mice was largely suppressed in α5/2 carotids (Figure 3A). Staining for the leukocyte adhesion receptors and inflammatory markers VCAM1 and ICAM1 showed strong elevation in the endothelium of the right carotid in WT mice but only minor changes in α5/2 mice (Figure 3B,C). Staining for CD68 as a marker for monocyte/macrophages showed strong inflammatory cell recruitment to the right carotid from WT mice, but only small changes in α5/2 mice (Figure 3D). Finally, we stained these tissues for pS536 NF-ĸB to assess activation of this critical inflammatory transcription factor. TAC induced strong NF-κB activation in the right carotid from WT mice but no significant changes in α5/2 mice (Figure 3E). Left carotids from both genotypes showed only minor changes in any of these markers. Together, these data show that inflammation and structural remodeling in this model of acute hypertension are drastically reduced in α5/2 chimera mice, consistent with a role of integrin α5

**Figure 2:**
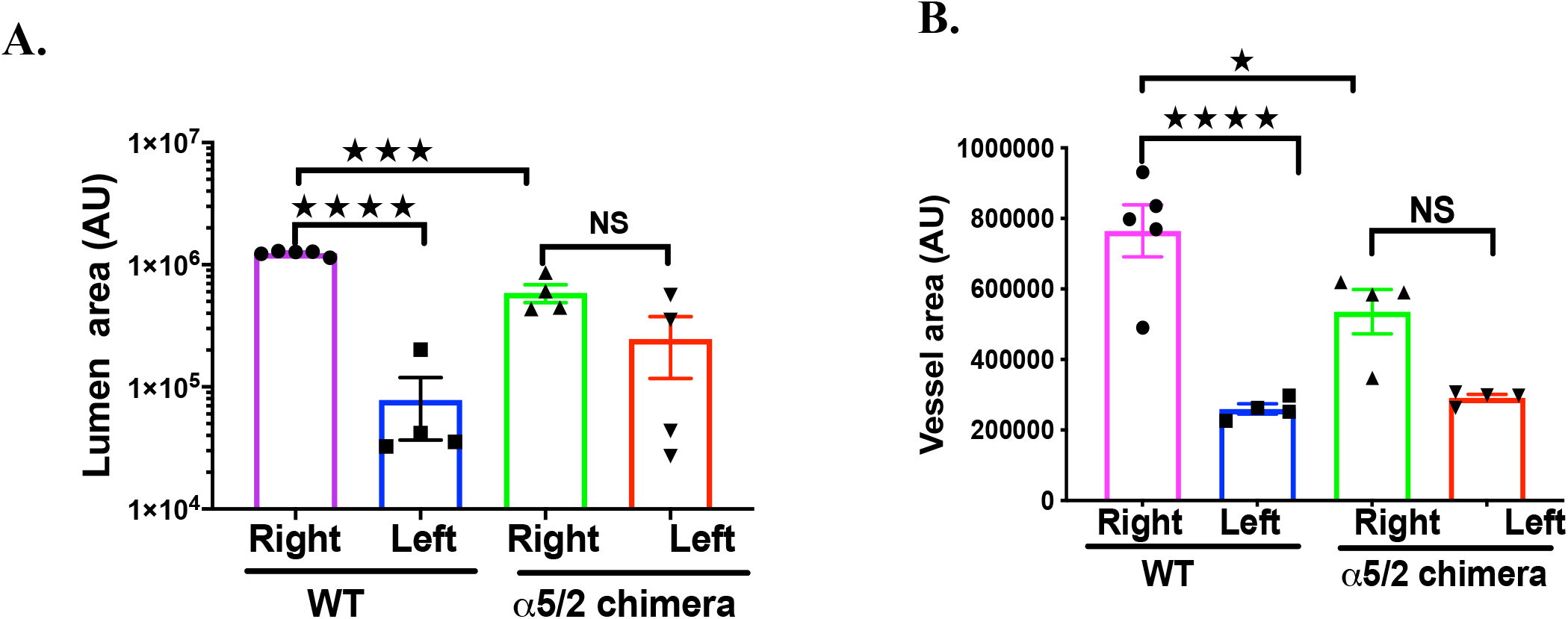
Transverse aortic constriction-induced artery remodeling in α5/2 mice. WT and α5/2 knock◻in mice at 1 week after TAC surgery. **A & B**. Vessel area and lumen diameter in carotid arteries from WT vs. α5/2 knock◻in mice. WT mice, n=5; α5/2 mice, n=4; Statistical analysis: one-way ANOVA with Tukey’s post hoc analysis; Values are means ± SEM; *p<0.05, ***p<0.001, ****p<0.0001 compared with WT mice.

**Figure 3:**
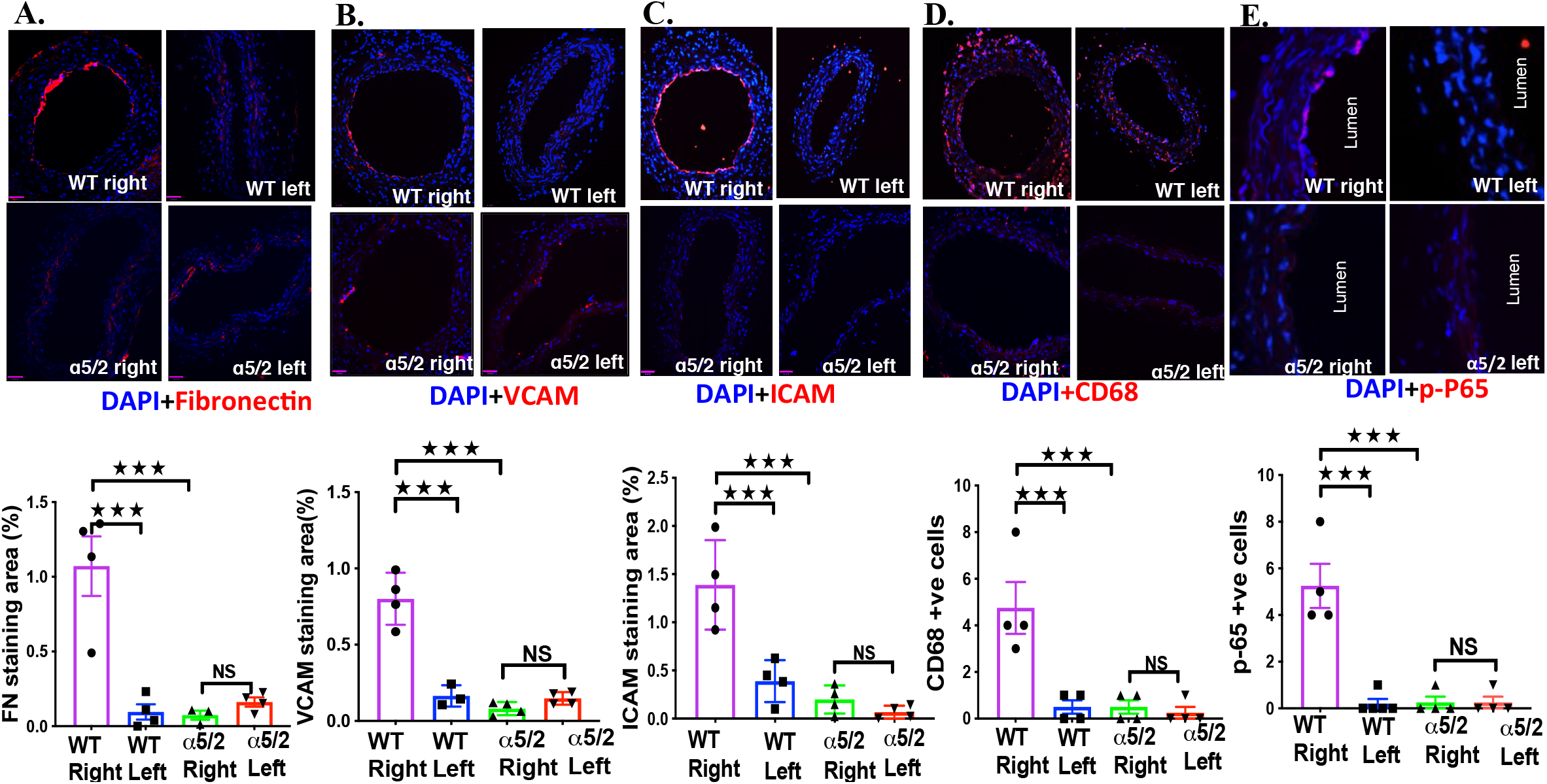
Transverse aortic constriction-induced fibronectin deposition and endothelial inflammation. WT and α5/2 knock◻in mice were subjected to TAC surgery. One-week post-surgery, carotid artery sections were stained for. **A**. Fibronectin; **B**. VCAM; **C**. ICAM; **D**. CD68; **E**. NFĸB p-65. Nuclei are counter stained with DAPI (blue). WT mice, n=4; α5/2 knock-in mice, n=4; Statistical analysis: one-way ANOVA with Tukey’s post hoc analysis; values are means ± SEM; ***p<0.001 compared with WT mice.

### Partial carotid ligation

Partial carotid ligation in mice decreases blood flow magnitude and introduces disturbances into the flow patterns in the common carotid, resulting in inflammatory activation of the endothelium and reduced lumen volume ^17, 18^. We first subjected WT mice to PCL surgery and examined carotid arteries at one-week post-surgery (Figure 4A). PCL resulted in reduced lumen diameter of the left carotid compared to the right carotid (Figure 4B). We also observed a dramatic increase in FN staining in the endothelial layer of the left but not right carotid arteries (Figure 4C). These observations prompted us to test the involvement of integrin α5 in these events by performing PCL in α5/2 mice. In contrast to WT mice, integrin α5/2 mice showed no change in the lumen area (Figure 4B). Further, ICAM1 and VCAM1 were strongly induced in the left carotid in WT mice but changed little in α5/2 mice (Figure 5A, B). Consistent with this finding, inflammatory cell recruitment marked by CD45 and CD68 was also attenuated in the left carotid of α5/2 mice (Fig. 5C.D) as was activation of NF-κB (Figure 5E). The contralateral right carotids of WT and α5/2 knock-in mice remained negative. PCL-induced artery inflammation and structural remodeling thus depend strongly on FN signaling through integrin α5.

**Figure 4.**
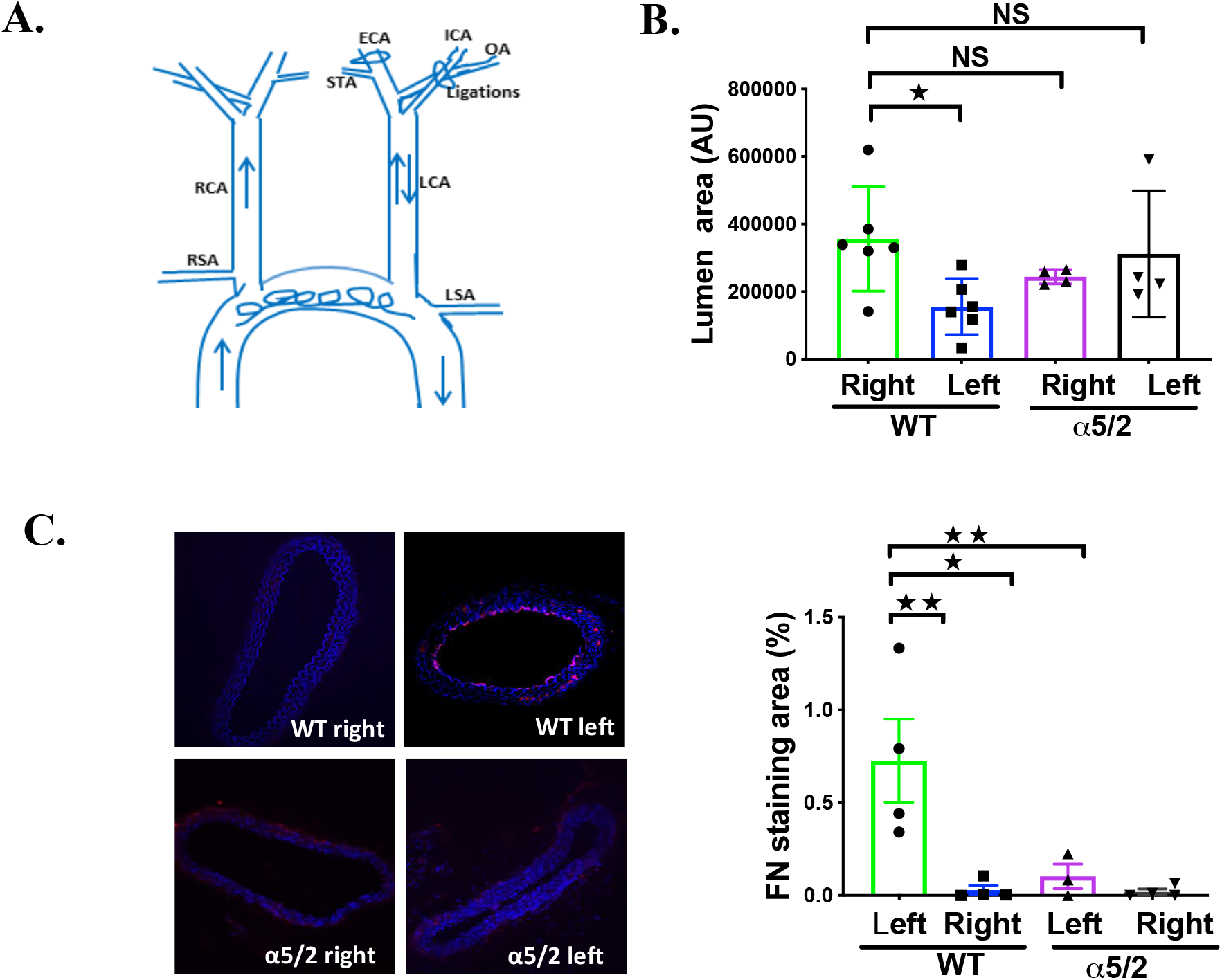
Partial carotid ligation-induced artery remodeling. **A.** WT and α5/2 knock◻in mice subjected to PCL surgery. Briefly, 3 out of 4 branches of the left common carotid artery (left external carotid, internal carotid, and occipital artery) were ligated with suture, while superior thyroid artery was left intact. **B**. One-week post-surgery, carotid arteries lumen diameter measured. Left carotid artery from WT mice have reduced lumen diameter compared to right carotid arteries form WT mice. In contrast to WT mice, integrin α5/2 mice showed no change in the lumen area. B. Increase in FN staining in WT mice left carotid compared with α5/2 mice. Nuclei are counter stained with DAPI (blue). ( WT mice, n=6; α5/2 knock-in mice, n=4); Statistical analysis: one-way ANOVA with Tukey’s post hoc analysis; Mean ± SEM ((*p<0.05, **p<0.01 compared with WT mice)

**Figure 5:**
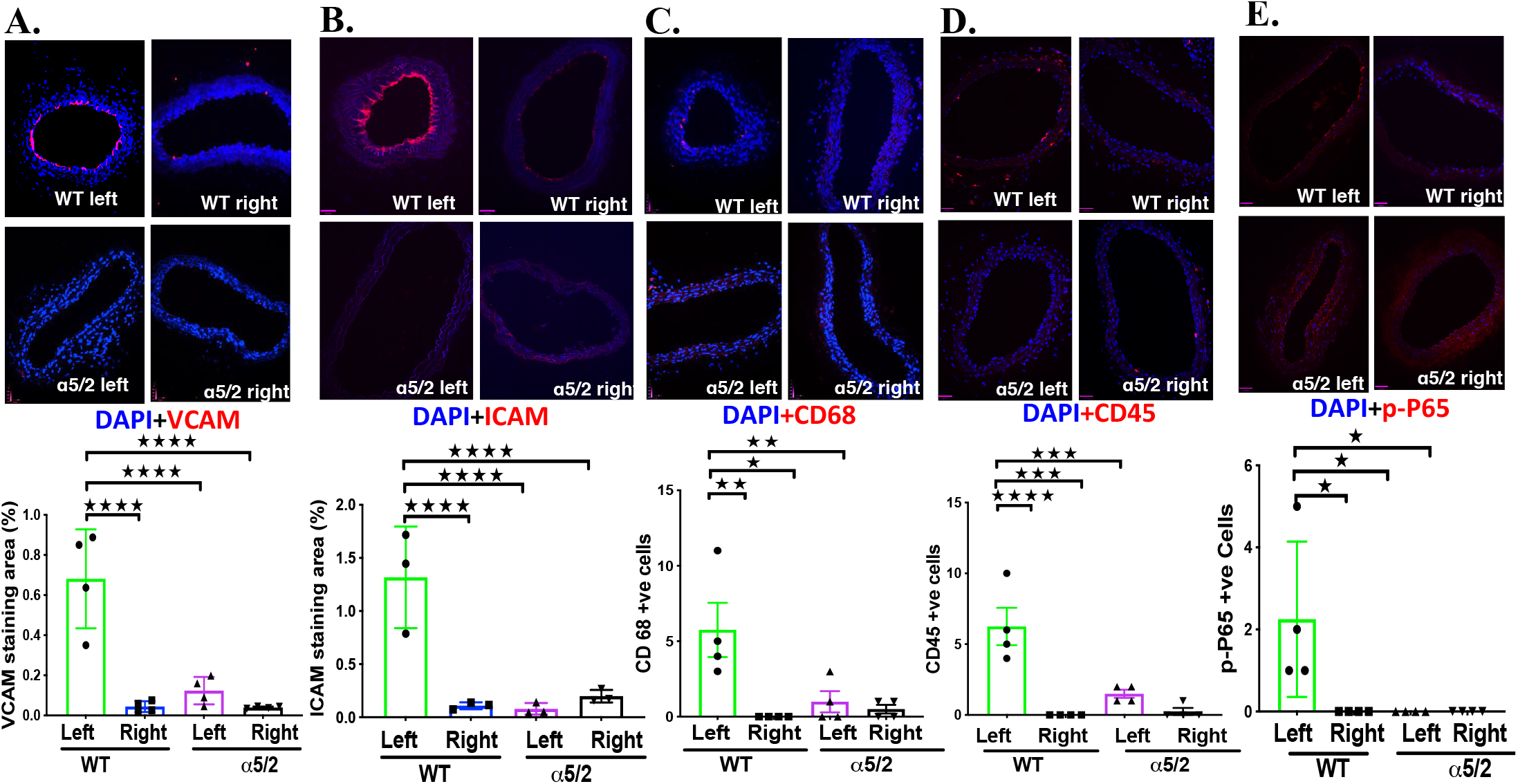
Partial carotid ligation-induced endothelial inflammation. WT and α5/2 knock◻in mice subjected PCL surgery. One-week post-surgery, carotid artery sections stained with following antibodies. **A**.VCAM; **B**. ICAM; **C**. CD68; **D**. CD45; **E**. NFĸB p-65. Nuclei are counter stained with DAPI (blue). ( WT mice, n=4; α5/2 knock-in mice, n=4); Statistical analysis: one-way ANOVA with Tukey’s post hoc analysis; Mean ± SEM ((*p<0.05, **p<0.01 compared with WT mice)

### Acute atherosclerosis after PCL in hypercholesterolemic ApoE^−/−^mice

PCL in hypercholesterolemic mice is used to model acute disturbed shear-induced atherosclerosis ^19, 20^. Lesions develop within a few weeks due to the very low and strongly oscillatory shear stress induced by surgery. To test the role of integrin α5 in this model, we crossed heterozygous α5/2 knock◻in mice with hypercholesterolemic ApoE^−/−^ mice (Jackson laboratories, B6.129p2-Apoetm1Unc/j; Cat#002052) to generated WT;ApoE−/− and α5/2;ApoE^−/−^ mice. Mice at 8-10 weeks were subject to PCL and fed a Western diet (RD Western diet #D12079B, Open Source Diet) for 3 weeks. H&E staining demonstrated significant atherosclerosis in the LCA of WT; ApoE^−/−^ mice whereas plaque in α5/2;ApoE^−/−^ mice was barely detected (Figure 6A). Staining lipids with Oil Red O confirmed the reduction in plaque in α5/2 mice (Figure 6B). Monocytes/macrophage (CD68^+^ cells) infiltration in the LCA was also markedly reduced in α5/2;ApoE^−/−^ mice (Figure 6C). As before, the contralateral side was unaffected and showed negligible plaque formation on this time scale. Plasma lipid profiles of total cholesterol, LDL cholesterol and triglycerides were not statistically different between WT and α5/2 mice (Figure 7). Overall, these results revealed that the α5/2 knock◻in strongly attenuated flow-dependent acute atherosclerosis.

**Figure 6:**
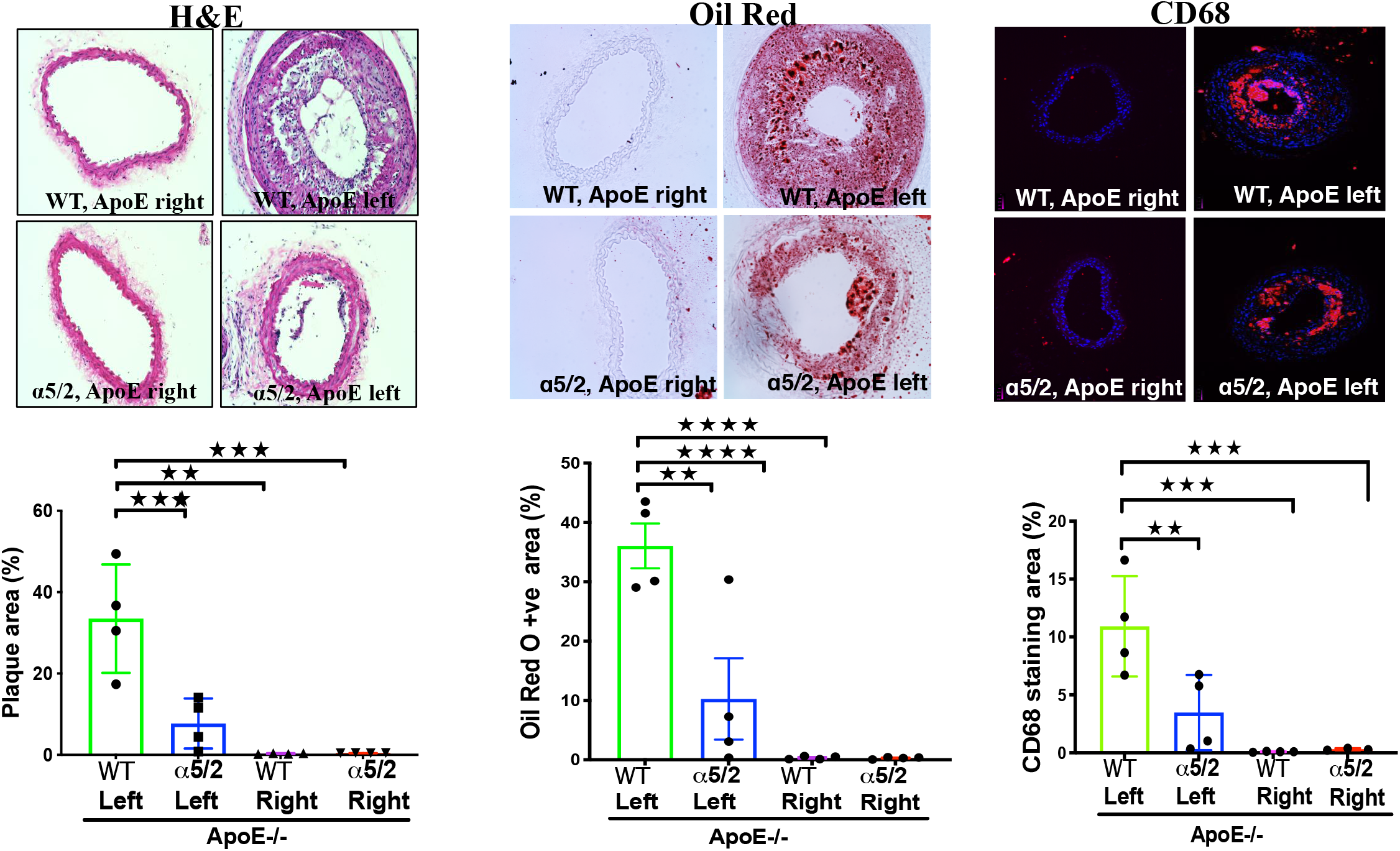
Partial carotid ligation in hypercholesterolemic mice. 8-10 weeks old WT and integrin α5/2 mice on the ApoE^−/−^ background were subjected to partial carotid ligation surgery.After the surgery, mice were fed a Western diet for 3 weeks and carotid arteries examined. **A**. H&E staining. **B**. Oil Red O staining. **C**. CD68 staining. WT;ApoE−/− mice, n=4; α5/2;ApoE−/− mice, n=4; Statistical analysis: one-way ANOVA with Tukey’s post hoc analysis; Values are means ± SEM; *p<0.05, **p<0.01 compared with WT mice.

**Figure 7:**
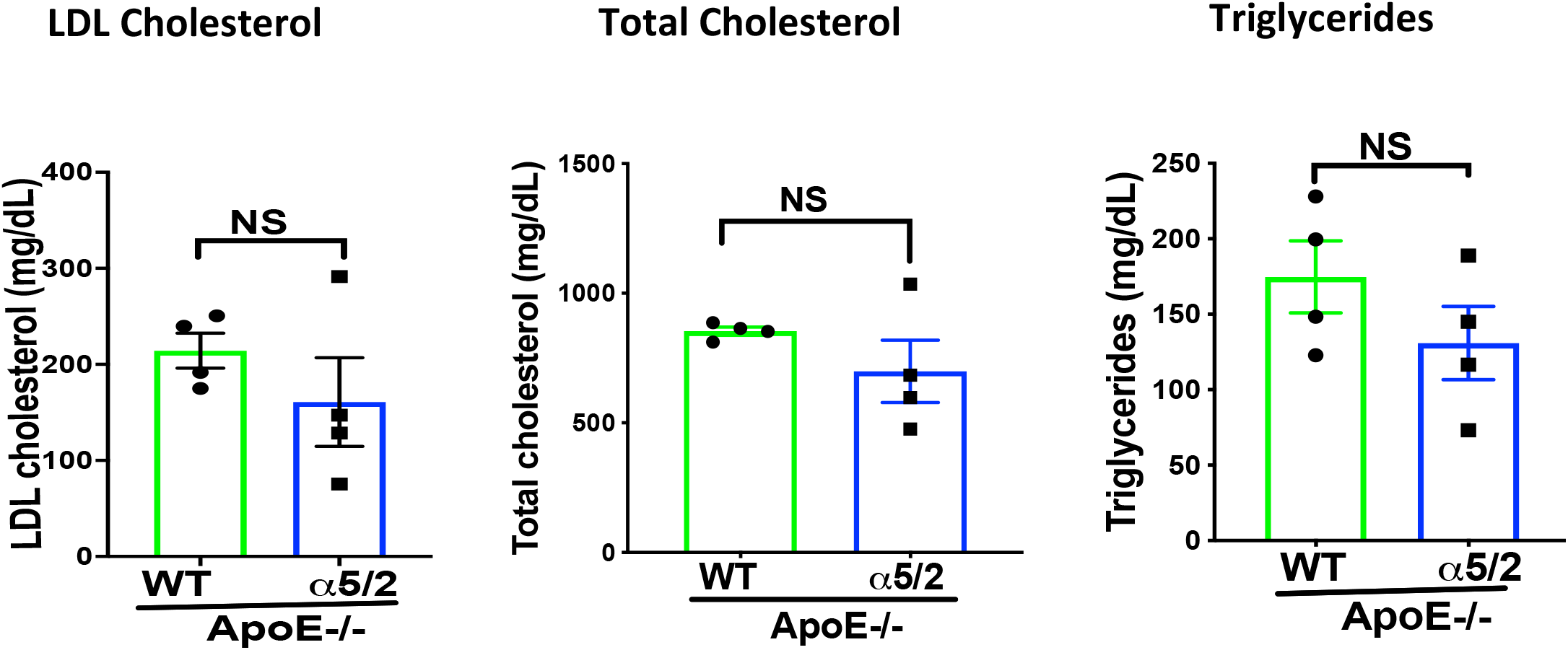
Plasma lipids. Plasma lipid profiles in WT;ApoE−/− and α5/2;ApoE−/− mice: **A.** Plasma LDL cholesterol; B. total cholesterol; C. Triglycerides. No significant differences were observed.

## Discussion

In this study, we report that the integrin α5/2 mutation in mice. which abolishes the inflammatory effects of FN, strongly affects vascular remodeling. In the TAC model of acute hypertension in the right carotid artery, α5/2 mice show markedly less FN deposition, less inflammatory activation of the endothelium, and less artery wall thickening. It should be noted that C57Bl/6 mice show excessive adventitial thickening in hypertension, perhaps related to the susceptibility of this strain to inflammatory stimuli ^21, 22^. These responses may thus be regarded as partially pathological in that remodeling overshoots the homeostatic goal. This effect was abolished by the α5/2 mutation, consistent with its ability to limit inflammation.

Partial carotid ligation is principally a model of flow-induced remodeling due to induction of low and oscillatory flow in the affected common carotid artery. Altered flow leads to inflammatory activation of the endothelium and inward remodeling to reduce lumen diameter, sometimes to near closure, a feature more in keeping with pathological than physiological processes, likely due to the oscillatory flow component. Consistent with high FN deposition in the endothelial layer of WT mice, inflammatory activation and inward remodeling were strongly blunted in α5/2 mice. To expand on these findings, we then examined effects of PCL in hypercholesterolemic mice, a model of acute flow-induced atherosclerotic plaque formation. Plaque size, lipid accumulation and inflammatory cell recruitment were all greatly decreased. These results strongly support a major role for inflammatory FN signaling in disturbed flow-induced vessel pathology.

Vascular remodeling in response to altered mechanical forces derived from blood flow and pressure is physiological when key variables such as tensional wall stress or fluid shear stress are restored to close to their initial levels ^2, 3^. Physiological remodeling requires inflammation and participation of immune cells but is carefully regulated and resolves when key variables return to their initial value or set point. By contrast, remodeling is pathological when initial set points are not restored due to either insufficiency or, more often, to overcompensation. For example, artery walls in hypertension thicken too much and lose elasticity, which exacerbates hypertension and its consequences for the microvasculature. Importantly, excessive or uncontrolled inflammation is the principle cause of pathological remodeling, typified by formation of atherosclerotic lesions in regions of low/disturbed shear stress ^23^. Adventitial fibrosis and stiffening in hypertension provides another example ^22^.

The current results identify FN and integrin α5 signaling as key elements of pathological vascular remodeling in acute models of both hypertension and disturbed flow. We note that mice in these experiments were total knock-ins, thus many cell types may contribute to the observed effects. Assessing effects in cell-type specific conditional knock-ins is an important direction for future work. While the ultimate goal of these studies is the identification of therapeutic targets, FN or integrin α5 themselves are essential for physiological processes and thus seem unlikely candidates for direct intervention. However, elements of downstream pathways have been discovered, including the interaction of phosphodiesterase 4D (PDE4D) with the α5 cytoplasmic domain and the interaction of PDE4D with the B55 subunit of protein phosphatase 2A ^24, 25^. Blocking these downstream pathways may be more promising avenues for translation, as are further investigations into mechanisms of α5 inflammatory signaling in atherosclerosis and hypertension.

## Methods

### Partial carotid ligation model

10-12 weeks old C57BL6, α5/2 integrin knock-in mice and ApoE^−/−^ mice were used for all animal studies. All the mice experiments were performed according to the approved protocol of the IACUC at Yale University. Mice were anaesthetized with ketamine and xylazine and surgery was performed. Briefly, 3 out of 4 branches of the left common carotid artery (left external carotid, internal carotid, and occipital artery) were ligated with suture, while superior thyroid artery was left intact. Mice sacrificed after isoflurane inhalation, perfused and vascular tree isolated along with carotid arteries at the indicated time points.

### Transverse aortic constriction

10-12 weeks old mice anaesthetized with ketamine and xylazine and hair was removed. A Para median incision was introduced with 1 mm sternum and each rib was cut individually, to expose heart and thymus, which was retracted to expose the aorta and carotid arteries. A curved 22-gauze needle with 9-0 sutures passed under aorta between carotid arteries and 27-gauze needle, space was placed on the arch prior to securing the ligature. The space was then removed, thus leaving a desired stenosis. The chest was closed with 3 to 5 simple interrupted sutures (5.0) followed by skin closure. Buprenorphine was given subcutaneously at 8 h interval following surgery. Mice sacrificed after isoflurane inhalation, perfused, and vascular tree isolated along with carotid arteries at indicated times.

### Immunofluorescence and histochemistry

Carotid arteries embedded in optimal cutting temperature, frozen on dry ice and stored at −80° C. Carotid arteries were sectioned on cryostat to generate 10-15 m sections. Cryosections were fixed in acetone for 10 min at −20 °C, blocked in IHC Tek antibody diluent for 1 hour at room temperature, and incubated with indicated antibodies in IHC Tek antibody diluent buffer. Antibodies were phosphor-NFĸβ-P65; FN (1:400; Sigma); VCAM1 (1:200 BD Bioscience); ICAM1 (1:200, Biolegend); CD68 (1:200, Abcam); CD45 (1:200, BD Biosciences); Sections were washed 3 times with PBS and incubated with Alexa flour 598-conjugated donkey anti rabbit or rat secondary antibody for 1 hour at room temperature. Slides were washed with PBS and mounted in Vectashield with DAPI. Images were acquired using Nikon 4 laser confocal microscope.

### Quantification and Statistical Analysis

NIH Image J program was used for quantification. Lumen diameter, vessel area, staining area was calculated from carotid artery sections. All graphs were created using GraphPad Prism software (San Diego, CA). Statistical analysis were performed using a one-way ANOVA with Turkeys post hoc analysis; p<0.05 was considered statistically significant. All data are expressed as mean ± SEM

## Acknowledgements

Lipid analysis was done by the Yale Mouse Phenotypic Center. This work was supported by National Institute of Health (RO1 HL75092) to MAS.

## Author’s contribution

MB designed the study, performed all experiments, and analyzed data and wrote the article. MB performed most of the studies at Yale and completed the analysis at University of Texas Health Sciences, San Antonio. JZ performed the mouse surgeries. MAS initiated the project, co-wrote the article, and provided financial support.

